# On the influence of the vascular architecture on Gradient Echo and Spin Echo BOLD fMRI signals across cortical depth: a simulation approach based on realistic 3D vascular networks

**DOI:** 10.1101/2024.05.30.596593

**Authors:** Mario Gilberto Báez-Yáñez, Jeroen C.W. Siero, Vanja Curcic, Matthias J.P. van Osch, Natalia Petridou

## Abstract

GE-BOLD contrast stands out as the predominant technique in functional MRI experiments for its high sensitivity and straightforward implementation. GE-BOLD exhibits rather similar sensitivity to vessels independent of their size at submillimeter resolution studies like those examining cortical columns and laminae. However, the presence of nonspecific macrovascular contributions poses a challenge to accurately isolate neuronal activity. SE-BOLD increases specificity towards small vessels, thereby enhancing its specificity to neuronal activity, due to the effective suppression of extravascular contributions caused by macrovessels with its refocusing pulse. However, even SE-BOLD measurements may not completely remove these macrovascular contributions. By simulating hemodynamic signals across cortical depth, we gain insights into vascular contributions to the laminar BOLD signal. In this study, we employed four realistic 3D vascular models to simulate oxygen saturation states in various vascular compartments, aiming to characterize both intravascular and extravascular contributions to GE and SE signals, and corresponding BOLD signal changes, across cortical depth at 7T. Simulations suggest that SE-BOLD cannot completely reduce the macrovascular contribution near the pial surface. Simulations also show that both the specificity and signal amplitude of BOLD signals at 7T depend on the spatial arrangement of large vessels throughout cortical depth and on the pial surface.

## 1. INTRODUCTION

Functional magnetic resonance imaging (fMRI) has revolutionized our understanding of brain function by allowing noninvasive mapping of brain activity. One of the most used fMRI techniques relies on the blood oxygen level-dependent (BOLD) signal, which is based on the combined effects of changes in local cerebral blood flow (CBF), cerebral blood volume (CBV) and oxygen metabolism (CMRO_2_) upon neuronal activation^1–3^. Due to recent advances in hardware, such as ultra-high magnetic field scanners (≥7T) and MR data acquisition strategies, the BOLD imaging technique enables the study of brain function at a high level of detail, i.e. at the mesoscopic organization of the cortex. As a consequence, it is now possible to measure BOLD fMRI activation as a function of cortical depth in human cortex and study neuronal activity across different cortical layers^4–17^. These breakthroughs underscore the potential of high-resolution fMRI as a valuable tool for investigating the fundamental processing within cortical micro-circuits and their intricate interactions.

While laminar BOLD fMRI holds great promise for advancing our understanding of cortical function, several challenges remain. Acquiring high-resolution BOLD fMRI data requires careful optimization of imaging parameters to balance spatial resolution, sampling rate, signal-to-noise ratio, and coverage. Laminar fMRI analysis techniques are still being refined, and validation against invasive techniques is necessary to confirm accuracy^18^. Moreover, the BOLD fMRI signal is an indirect measurement of neuronal functioning. The BOLD fMRI signal is a mixture of effects related to hemodynamic changes induced by neurovascular coupling, the vascular architecture within the sampled volume, and the biophysical interaction of oxygenated blood and tissue. Because BOLD fMRI measures neuronal activity through hemodynamics, its ultimate resolution relies on the spatial extent of neuronal-evoked hemodynamic changes, along with how these changes evolve over time^19–21^.

Numerous laminar fMRI studies, with spatial resolutions reaching ≤1mm in all directions, have examined the spatial characteristics of the recorded hemodynamic signals across cortical depth, to enhance our understanding and refine spatial specificity of laminar fMRI^7,13,22–27,57^. Gradient Echo (GE)-BOLD is the most commonly used technique to measure brain activation across cortical depth. Nevertheless, GE-BOLD is sensitive to both macro- and microvascular signal contributions, which impact the specificity of the underlying neuronal activity, especially towards the pial surface. Computational simulations based on mono-sized randomly oriented cylinder models have shown that Spin Echo (SE)-BOLD can minimize the extravascular contribution from large vessels^28–31^. Therefore, SE-BOLD increases the specificity towards the microvasculature and is thus considered to be more specific to the location of neuronal activity^32,33^. However, SE-BOLD fMRI measurements may not completely remove the macrovascular contribution – macrovessels may still influence the detected BOLD signal, especially near the pial surface^23,34^ due to several possible factors, such as the impact of the echo planar imaging (EPI) readout^35,36^, the dependence on the echo-time (TE) selection^37^ or the different diffusion regimes between the cortical gray matter (GM) and the cerebrospinal fluid (CSF)^30,38^. For that reason, it is necessary to investigate the GE-BOLD and SE-BOLD signal formation across cortical depth using computational simulations based on realistic 3D cortical vascular networks.

In this work, we used realistic 3D vascular models of mice obtained by two-photon microscopy techniques, with a modified artery-vein ratio in order to mimic human cortical vasculature features, to simulate specific oxygen saturation states and biophysical interactions to characterize the intravascular and extravascular signal contribution of the vascular architectures to the GE-BOLD and SE-BOLD signals across cortical depth. This computational approach can help to understand the influence of the vascular architecture on the GE-BOLD and SE-BOLD signal formation across cortical depth, and the impact of the pulse sequence parameters on the BOLD signal changes at submillimeter acquisitions.

## 2. MATERIAL AND METHODS

### 2.1 Generation of realistic 3D vascular models based on two-photon microscopy data

We utilized four realistic 3D vascular models acquired with two-photon microscopy techniques from the parietal cortex of mice as obtained by Blinder et. al. [2013]^39^ (***Figure 1***). Using an in-house MATLAB script, we reconstructed the vasculature using the spatial coordinates of the vessel segments and their associated diameter information. The vascular data, as provided by Blinder et. al. [2013]^39^, was stored in a vectorial structure – where spatial positions comprised an *m*-by-3 vector, with *m* representing the number of vessels of the vascular model, and two *m*-by-1 column vector representing the diameter and length of each vessel segment, respectively. Capillaries are effectively represented by their length and diameter due to their uniform structure. However, arteries and veins, with their intricate shapes, require subdivision into multiple segments to accurately capture their branching patterns, varying diameters, and curvature. More details on the vascular data can be found in Blinder et. al. [2013]^39^.

**Figure 1.**
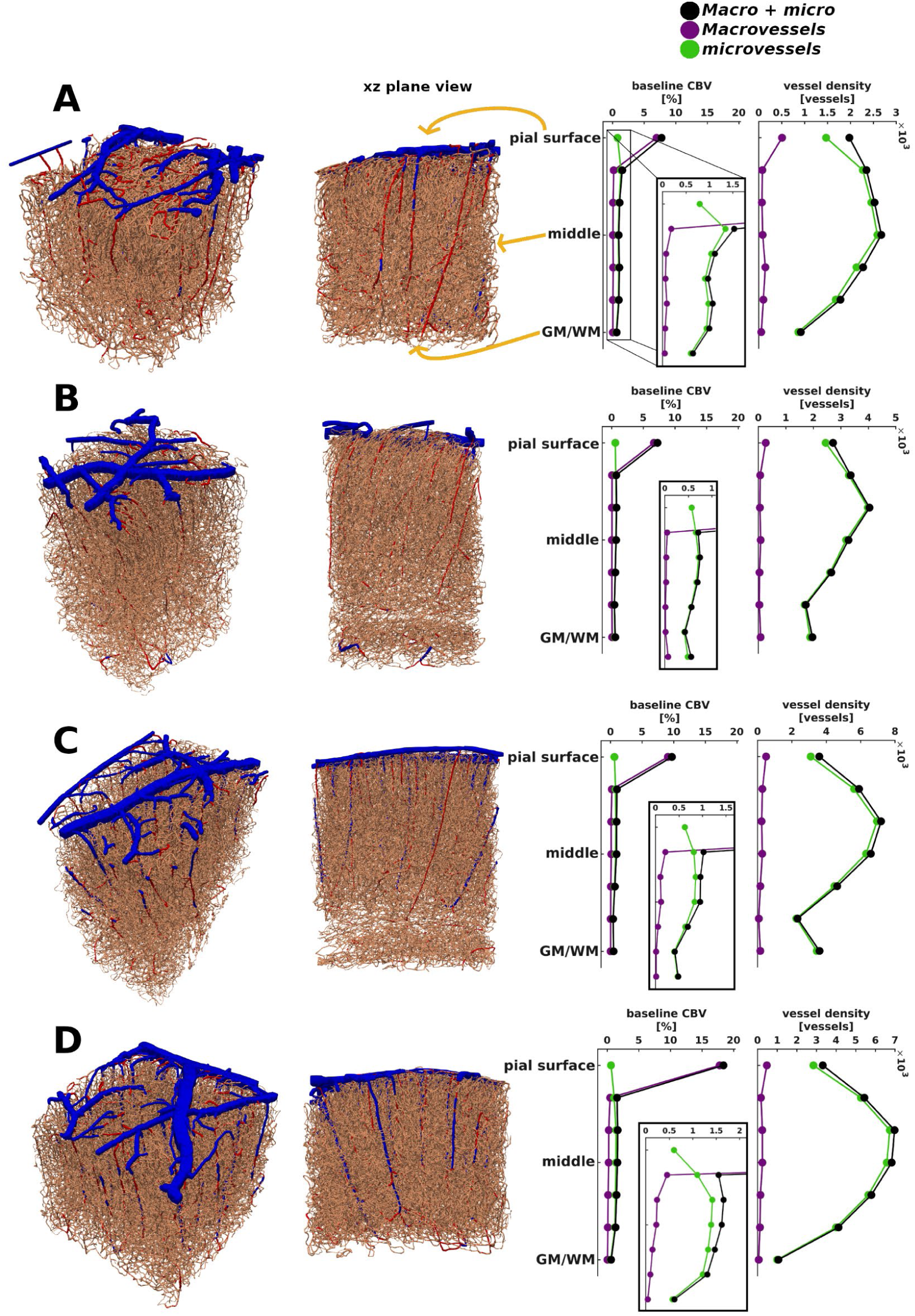
Four realistic 3D vascular models (A-D models) obtained by means of two-photon microscopy techniques^39^. The realistic vascular models are comprised of three vessel types – arteries in red and veins in blue (both considered thorough the manuscipt as macrovessels), and capillaries, small arterioles, and venules in beige (considered as microvessels). On the right hand side of each vascular model, baseline CBV and vessel density features are displayed for both macrovessels (purple dotted lines) and microvessels (green dotted lines) – and the total sum in black dotted lines. The models were subsequently used to compute the spatial distribution of frequency shifts assuming different oxygen saturation levels per vascular compartment, and from the frequency shifts the corresponding R2^(^*^)^ decay rate effects and the BOLD signal changes are obtained (sections 2.2-4).

In this work, vessels smaller than 6 micrometers in diameter were labeled as capillaries^33,40^. To mimic human brain characteristics, the remaining vessels were divided into a 3:1 artery-vein ratio based on diameter. This reflects the 3:1 ratio in humans^41^ compared to approximately 1:3 in mice^39,42,43^. Vessels with diameters ranging from 11 µm to 36 µm were designated as veins, constituting the smaller portion, while those with diameters from 6 µm to 11 µm were labeled as arteries^59,60^. Thus, the resulting 3D vascular models included arteries and veins (referred to as macrovessels from here onwards) along with capillaries, small arterioles, and venules (microvessels). The simulated voxel size generated by the vascular models is about 1 mm^3^ isotropic. We defined seven cortical layers evenly distributed between the cortical pial surface and the white matter/gray matter (WM/GM) boundaries (see ***Figure 1***). It’s important to note that these layers were not intended to represent histological laminae or to delineate any vascular features. The number of vessels was computed as the sum of the vessels lying within each of the defined cortical layers. Moreover, the baseline CBV of each cortical layer was computed as the sum of the volume of all its vessels. The volume of a single vessel was calculated from the vessel diameter and length assuming a cylindrical shape.

### 2.2 Implementation of oxygen saturation levels for each vascular compartment

We simulated different oxygen saturation levels per vascular compartment^53^, which were maintained constant over time, i.e., steady-state oxygen saturation levels were assumed. The baseline oxygen saturation (SO_2_) values used in each vascular compartment were dependent on the oxygen saturation imposed on the veins, as follows:

- SO_2_ in arteries (SO_2_art) = 95%;
- SO_2_ in capillaries = SO_2_art - ((SO_2_art - SO_2_vein) / 2);
- SO_2_ in veins (SO_2_vein) = [50%, 54%, 59%, 64%, 68%, 73%, 78%, 82%, 87%, 92%].

We extended the simulation oxygen saturation range beyond physiological plausible values (interval that ranges from oxygen saturation values of 60% in the resting state to 80% in the active state), in order to gain a broader understanding of the BOLD signal formation process.

### 2.3 Simulation of the MR signal accounting for intravascular and extravascular signal contributions

BOLD fMRI signals include extravascular and intravascular components, influenced, mainly, by factors like cerebral blood volume and oxygen saturation. The total MR signal was calculated by summing the extravascular signal with both arterial and venous intravascular contributions:

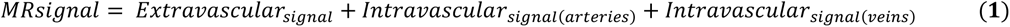

Simulations shown here were computed for gradient echo and spin echo at 7T with an angular orientation of the normal vector arising from the cortical pial surface of the realistic vascular model parallel to the main magnetic field.

#### 2.3.1 Simulation of the arterial and venous intravascular signal contribution

In this study, we assumed the intravascular contribution (*R*2^(∗)^ *_dHb_*= *R*2(∗)0,*in* + *R*2*_SO_*_2_) to the BOLD signal to be non-zero for the arterial and venous compartment. This decision was based on the observation that, at high magnetic fields, the contribution of the arterial and venous compartment tends to be significant at specific oxygen saturation levels^32,33^. On the other hand, we assumed an intravascular component in the microvascular compartment to be null, given that the *R*2^(∗)^*_dHb_* of the capillaries has not been well-characterized due to the high heterogeneity in hematocrit levels across the cortical depth^55^ and oxygen saturation levels across the capillary bed.

Therefore, we implemented the intravascular arterial and venous contribution for SE as 1/*R*2_0,*in*_ = *T*2_0,*in*_ (≈ 53 ms)^34^, and *R*2*_SO2_* component dependent on oxygen saturation level using the quadratic relation as defined by Uludag et al., [2009]^33^, weighted by the corresponding blood volume fraction (see ***Figure 1***). Moreover, we implemented the intravascular arterial and venous contribution for GE as 1/*R*2^∗^_0,*in*_ = *T*2^∗^_0,*in*_ (≈ 10 ms)^34^, and *R*2*_SO2_* component dependent on oxygen saturation level using the relations defined by Uludag et al., [2009]^33^, weighted by the corresponding blood volume fraction (see ***Figure 1***).

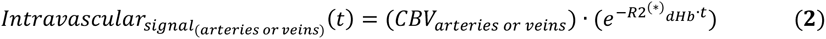

Although the intravascular decay rate for both GE and SE is influenced by hematocrit level, we assumed a constant value of hematocrit across vascular compartments in our simulations. This decision aimed to reduce one degree of freedom in the simulations.

#### 2.3.2 Simulation of the extravascular signal contribution

The extravascular *MRsignal* was computed by modelling the interaction of moving spins within the local magnetic field distortions induced by the different oxygen saturation levels^29^ (section 2.2). We computed local frequency shifts created by the SO_2_ levels of both the macro- and microvascular compartments (***Figure 2***).

**Figure 2.**
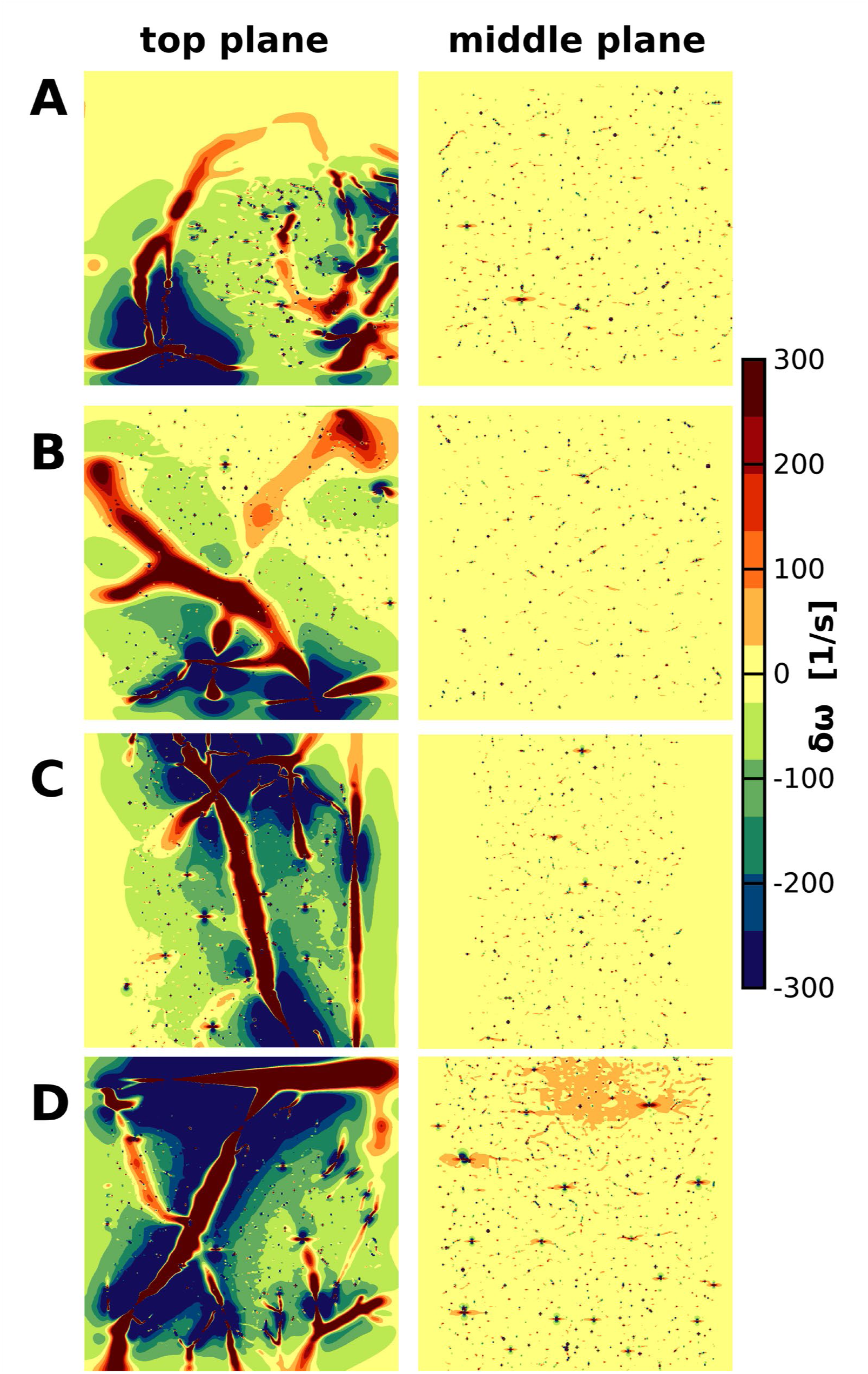
Local frequency field distortions produced by the realistic vascular models for an exemplary oxygen saturation level (SO_2vein_ = 78%). The frequency field maps are shown for two different locations inside the simulated voxel: the top plane, situated approximately 20 μm in depth from the cortical pial surface, and the middle plane, positioned around 500 μm deep relative to the cortical pial surface. The main magnetic field was simulated to be orthogonal to the cortical pial surface and thus the depicted planes. The unique macrovascular structure of the four vascular models at the pial surface level (top plane) produces a distinctive local spatially inhomogeneous frequency field signature. However, at the middle plane, the vascular architecture is primarily composed of small vessels, leading to a relatively similar degree of induced inhomogeneous frequency field across the models.

The local frequency shift caused by a vessel segment was modeled as the dipolar response of a finite cylinder, presuming negligible effects on the cylinder extremities^44,45^. The local frequency shift *δω*(*r*) in [1/s] for each vessel segment was computed using:

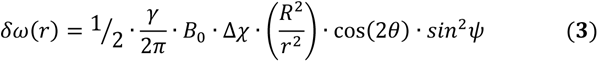

where γ is the hydrogen gyromagnetic ratio = 267.5E6 [rad/(s·T)], *B*_0_ is the main magnetic field (7 [T]), Δ*χ* = 4*π* · 0.276 *ppm* · *HcT* · (1 − *SO*_2_) [-] is the susceptibility difference produced by the *SO*_2_ in the vessel/cylinder^46^ and the hematocrit level *HcT* (= 0.45 [-]), *R* is the vessel radius in [µm], *r* is the Euclidean distance from the center line of the cylinder to a particular spatial position in the simulation volume in [µm], *θ* is the angle between the cylinder and the spatial position in [rad], and *ψ* is the angle between the orientation of the cylinder and the main magnetic field in [rad].

The dephasing experienced by a bulk of diffusing spins *N_spins_* was simulated using a Monte Carlo approach (20 repetitions with 5·10^7^ spins each repetition). An ensemble of spins senses the extravascular local frequency shifts with a diffusion coefficient of D = 1 [µm^2^/ms], while assuming isotropic diffusion^29^. The calculation of the spin dephasing was obtained through,

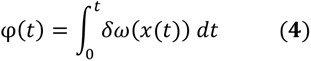

where φ(*t*) is the phase acquired during the simulation time *t* and *δω*(*x*(*t*)) is the local frequency shift at spin position *x* at each time-step *t*. The phasing experienced for each spin was stored across all simulation time-steps (time step = 0.025 ms). For SE sequences, the acquired phase during the echo time was multiplied by -1 (change in polarity) after TE/2, simulating the effect of the 180-degree refocusing radiofrequency pulse. Using equation **(5)** we can obtain the normalized extravascular MR signal.

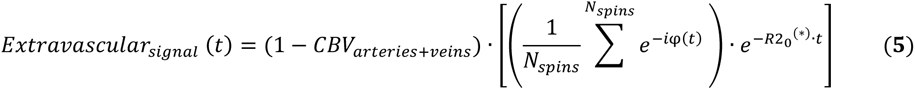

Where *R*2_0_^(∗)^ = 1/T2_0_^(^*^)^ is the intrinsic decay rate in cortical tissue, and R2’, expanded in the term inside the parenthesis, is the decay rate induced by the interaction of the diffusing spins in a local inhomogeneous frequency field. We used the intrinsic tissue T2_0_* (≈ 28 ms) relaxation time for GE and the intrinsic tissue T2_0_ (≈ 50 ms) relaxation time for SE according to the nonlinear relationship given by Khajehim et. al. [2017]^47^ for cortical gray matter at 7T (see ***Table 1***).

**Table 1.** Biophysical and pulse sequence parameters used to compute a BOLD signal response.

| biophysical and MR pulse sequence parameters |  |
| --- | --- |
| diffusion coefficient (extravascular tissue) | 1 $\mu\text{m}^2/\text{ms}$; isotropic motion |
| static magnetic field strength | 7 tesla |
| hematocrit (HcT) | 0.45 |
| echo-time (TE)<br>(used in **Figure 3 and 4**) | gradient echo = 27 ms<br>spin echo = 50 ms |
| TE range to estimate R2(*) and BOLD signal changes TE-dependency (used in **Figure 5 and 6**) | 5 ms to 80 ms, step of 5ms |
| time-step | 0.025 ms |
| number of spins | 20 Monte-Carlo repetitions with<br>5·10<sup>7</sup> spins each repetition |
| GE: T2<sub>0</sub>* cortical grey matter at 7T<br>SE: T2<sub>0</sub> cortical grey matter at 7T | 28 ms<br>50 ms |
| GE: intravascular arterial T2<sub>dHb</sub>* at 7T | 9 ms @ SO<sub>2</sub> = 95% |
| SE: intravascular arterial T2<sub>dHb</sub> at 7T | 49 ms @ SO<sub>2</sub> = 95% |
| GE: intravascular venous T2<sub>dHb</sub>* at 7T | [4, 5, 5.4, 6, 6.7, 7.4, 8, 8.7, 9.3, 9.7]<br>ms @ SO<sub>2</sub> = [50%, 54%, 59%, 64%,<br>68%, 73%, 78%, 82%, 87%, 92%] |
| SE: intravascular venous T2<sub>dHb</sub> at 7T | [5, 7, 8.2, 10.1, 12.6, 15.9, 20.6,<br>26.9, 35, 44.3] ms @ SO<sub>2</sub> = [50%,<br>54%, 59%, 64%, 68%, 73%, 78%,<br>82%, 87%, 92%] |

To confine spins within the simulation space, voxel boundary conditions were set to infinite space. Spins exiting the voxel re-entered the imaging volume on the opposite side, preserving their magnetization history. However, spins reaching the pial surface and WM/GM boundary were considered invalid iterations and reiterations were performed. Additionally, spin exchange between vascular compartments was prohibited, establishing an impermeable vascular network

### 2.4 GE R2* and GE BOLD signal change, and SE R2 and SE BOLD signal change across cortical depth

The *MRsignal* and the corresponding GE R2* and SE R2 were computed using their respective echo time as described in section 2.3. Given that the behavior of the *MRsignal* (***Eq. 1***) presents oscillations due to its multi-exponential nature, we simply approximate the R2^(^*^)^ decay rate value fitting a polynomial of degree one, i.e. a linear fit, on the natural logarithm of the *MRsignal*, 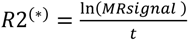, for GE and SE, respectively. The BOLD signal change in [%] was defined as the relative change using the 59% oxygen saturation state as the reference / baseline condition (SO_2vein_ = 59%), i.e.,

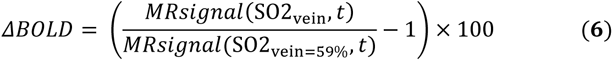

Hence, we conducted three different experiments/simulations:

1. To quantify the global dephasing effects of GE R2* and SE R2, we conducted Monte Carlo simulations at the voxel level across the various oxygen saturation states (section 2.2). Additionally, we calculated the GE R2*/SE R2 ratio as a proxy for vessel size signal contribution^29,48^. Furthermore, we calculated the impact of the extravascular and intravascular signal contribution for different oxygen saturation values for all vascular models.
2. Subsequently, we used Monte Carlo simulations to calculate the contributions of GE R2* and SE R2, and the corresponding BOLD signal changes, for seven cortical layers. As mentioned above, these cortical layers were evenly defined between the pial surface and the white matter/gray matter (WM/GM) boundaries (see ***Figure 1***). Additionally, we calculated the GE R2*/SE R2 ratio and the ((GE BOLD / SE BOLD) - 1) signal ratio as a proxy for vessel size signal contribution^29,48^.
3. Finally, we simulated the GE R2* and SE R2 dephasing rate across the seven cortical layers for different echo times ranging from TE = 5 ms to 80 ms with increments of 5 ms. Additionally, we examined the respective BOLD signal changes.

The computational pipeline was implemented using MATLAB and JULIA. The biophysical parameters used to compute the GE R2* and SE R2, and the respective BOLD signal changes are summarized in ***Table 1***.

### 2.5 Validation of computational pipeline through generation of the Boxerman^29^ plot

To validate the computational framework, we reproduced results from Boxerman et al. [1995]^29^ and Kiselev et al. [1999]^30^. Using mono-sized randomly oriented cylinders, we computed ΔR2’ effects for GE and SE at 1.5T with parameters from Kiselev et al. [1999]^30^. This is shown in ***Supplementary*** Figure 1. Additionally, we calculated GE BOLD and SE BOLD signal changes to understand BOLD contrast at 7 Tesla using this simplified model, detailed in ***Supplementary*** Figure 2.

## 3. RESULTS

*Figure 1* illustrates the four vascular models utilized in this study, encompassing arteries, microvessels (including capillaries, small arterioles/venules), and veins. While obtained from different mice within the same cortical region (parietal), the capillary beds in all four models demonstrate similar characteristics in vessel radius, baseline CBV, and vessel density across cortical depth, with somewhat comparable topologies. However, macrovascular architecture varies among the models, leading to distinct differences in large vessel organization.

The local inhomogeneous frequency shifts produced by the four realistic vascular models for an exemplary oxygen saturation level (SO2vein = 78%) are depicted in Figure 2. Two orthogonal planes are depicted: one approximately 20 μm deep from the cortical surface (top plane), and the other around 500 μm deep relative to the cortical pial surface (middle plane). Near the cortical surface, significant frequency field inhomogeneities arise from the macrovasculature, primarily pial veins, with minor contributions from smaller vessels. Conversely, microvessels dominate the middle planes, resulting in slight alterations to the frequency field homogeneity, affecting the behavior of moving spins. The distinct macrovascular architecture at the pial level creates specific local frequency field signatures, while the middle planes exhibit comparable degrees of induced inhomogeneity across the models.

*Figure 3* illustrates the global, i.e. averaged signals over the complete voxel, dephasing in terms of voxel’s R2^(^*^)^ decay rates using GE (TE = 27 ms) and SE readouts (TE = 50 ms) for the four vascular models. Each solid line in the graphs represents the mean value, while the shaded area represents the 95% confidence interval computed from the repeated Monte Carlo simulations. The grey shaded area covering the oxygen saturation range between 60% to 80% represents the plausible physiological values expected in experimental fMRI data. The horizontal dashed line in Figure 3a and ***3b*** represents the expected R2^(^*^)^ in the case where there are no susceptibility differences, i.e., the intrinsic R2_0_^(^*^)^ of tissue.

**Figure 3.**
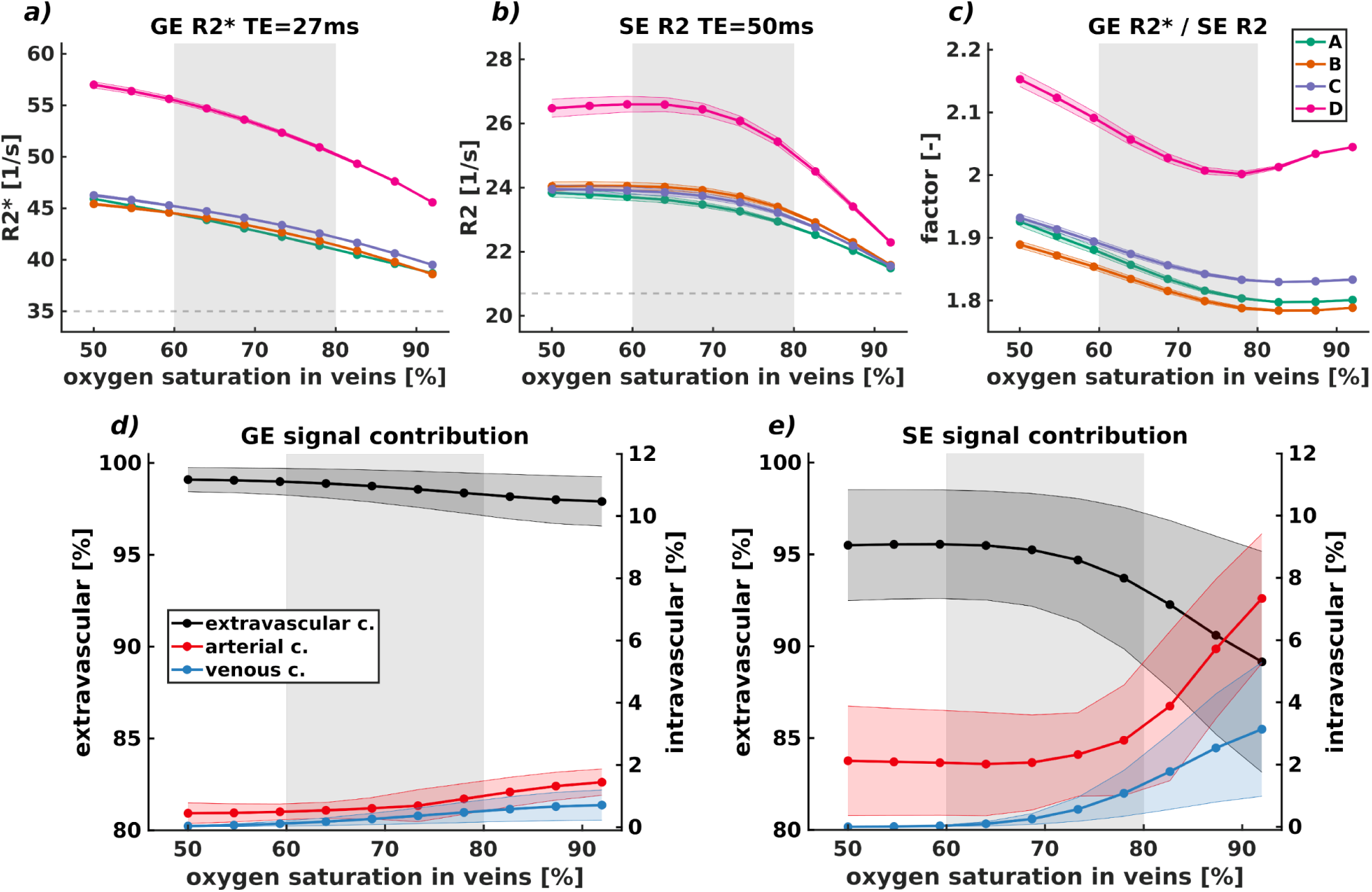
Global dephasing, characterized by voxel R2^(^*^)^ decay rates, and the extravascular and intravascular contributions to this simulated for GE (TE = 27 ms) and SE (TE = 50 ms) readouts across four vascular models and varying static oxygen saturation values at 7T. Each solid line in the results represents the mean value, while the shaded area in **(a)**, **(b)** and (**c)** indicates the 95% confidence interval computed through repeated Monte Carlo simulations, and in **(d)** and **(e)** the 95% confidence interval averaged across all vascular models. The grey shaded region, encompassing an oxygen saturation range between 60% to 80%, reflects the plausible physiological values anticipated in human fMRI data. The horizontal dashed line, in **a)** and **b)**, denotes the R2^(^*^)^ devoid of susceptibility differences corresponding to the intrinsic R2^(^*^)^ of tissue. GE R2* **(a)** shows a larger effect as compared to SE R2 **(b)**. The ratio of GE R2* and SE R2 **(c)** varies between 1.80 to 2.10 in the plausible physiological range. At low oxygen saturation levels (SO2vein = 50%), the range is approximately from 1.89 to 2.15, while at higher oxygen saturation levels (SO2vein = 92%), this ratio is approximately in the range from 1.79 to 2.05. The GE R2* and SE R2 decay rate behavior for the A, B, and C models displayed largely similar values, resulting in overlapping curves. Conversely, the D model featured much higher ratio values as compared to the other models. The contributions of extravascular, arterial, and venous intravascular components to GE and SE signals are depicted in **d)** and **e)**. The black line represents the behavior of the extravascular signal component. The arterial and venous intravascular signal contributions are illustrated by red and blue lines, respectively. In GE signal contribution, the dominant signal originates from the extravascular compartment (∼98% signal contribution), whereas both arterial and venous intravascular contributions are minimal. Conversely, in SE signal contribution, the intravascular signal, from both arterial and venous sources, exhibits a more significant contribution compared to GE. Depending on the oxygen saturation level, the arterial component can contribute up to 8% of the signal, while the venous component can contribute up to ∼4% and thus the extravascular compartment is reduced -constituting a signal contribution of about ∼90%-95% dependent on the oxygen saturation level.

All vascular models consistently exhibited a nonlinear response across various oxygen saturation levels (see Figure 3a and ***3b***). Increasing oxygen saturation within the voxel led to a nonlinear decrease in both GE R2* and SE R2 decay rates, with SE R2 showing a more pronounced impact compared to GE R2*. Additionally, the confidence interval for SE R2 was slightly larger for low oxygen saturation values compared to higher levels. This difference was not observed in GE R2*, where confidence intervals remained similar across all oxygen saturation levels. The ratio between GE R2* and SE R2 demonstrated a distinctive wave-like shape (see Figure 3c). The lowest point of the curve fell within the plausible physiological range, with values ranging from 1.79 to approximately 2.00, depending on the vascular model. At low oxygen saturation levels (SO2vein = 50%), the ratio ranged from 1.89 to 2.15, while at higher levels (SO2vein = 92%), it ranged from 1.79 to 2.05. Models A, B, and C exhibited similar GE R2* and SE R2 decay rate behaviors, resulting in overlapping and intersecting curves. In contrast, model D showed considerably higher ratio values compared to the other models.

The extravascular, arterial, and venous intravascular contributions to GE and SE signals are depicted in Figures 3d and ***3e***, with solid lines representing mean values and shaded areas indicating the 95% confidence interval from repeated Monte Carlo simulations across all vascular models. The grey shaded area encompasses oxygen saturation values between 60% to 80%, considered physiologically plausible. The black line represents the extravascular signal component, while the red and blue lines represent arterial and venous intravascular signal components, respectively.

In GE signal contribution, the dominant signal comes from the extravascular compartment (∼98% signal contribution), with minimal intravascular contributions from both arterial and venous sources, each less than 2%. Conversely, in the SE signal, intravascular signal contributions, particularly from arterial sources, are more substantial. Depending on the oxygen saturation level, the arterial component can contribute up to 8%, while the venous component can reach approximately 4%. Consequently, the extravascular compartment constitutes an average signal contribution of about 90% to 95%, depending on the oxygen saturation level.

In Figure 4, we illustrate the relationship between the R2^(^*^)^ decay rate (panels ***a)*** to ***c)***) and BOLD signal changes (panels ***d)*** to ***f)***) for both GE (TE = 27 ms) and SE (TE = 50 ms) readouts across seven equidistant layers of the cortex for all vascular models. The panels ***c)*** and ***f)*** present the resulting GE R2*/SE R2 ratio and the ((GE BOLD / SE BOLD) - 1) signal ratio, respectively. Each solid line in the results represents the mean value, while the shaded area represents the 95% confidence interval computed by the Monte Carlo simulation averaged across all the oxygen saturation levels – where the dashed line represents the boundary produced by the lowest oxygen saturation value and the dotted line represent the boundary given by the highest oxygen saturation value. The independent contribution for each oxygen saturation can be found in ***Supplementary*** Figures 3a and ***Supplementary*** Figure 3b. The horizontal dashed line in ***a)*** and ***b)*** represents the intrinsic R2_0_^(^*^)^ of tissue, respectively.

**Figure 4.**
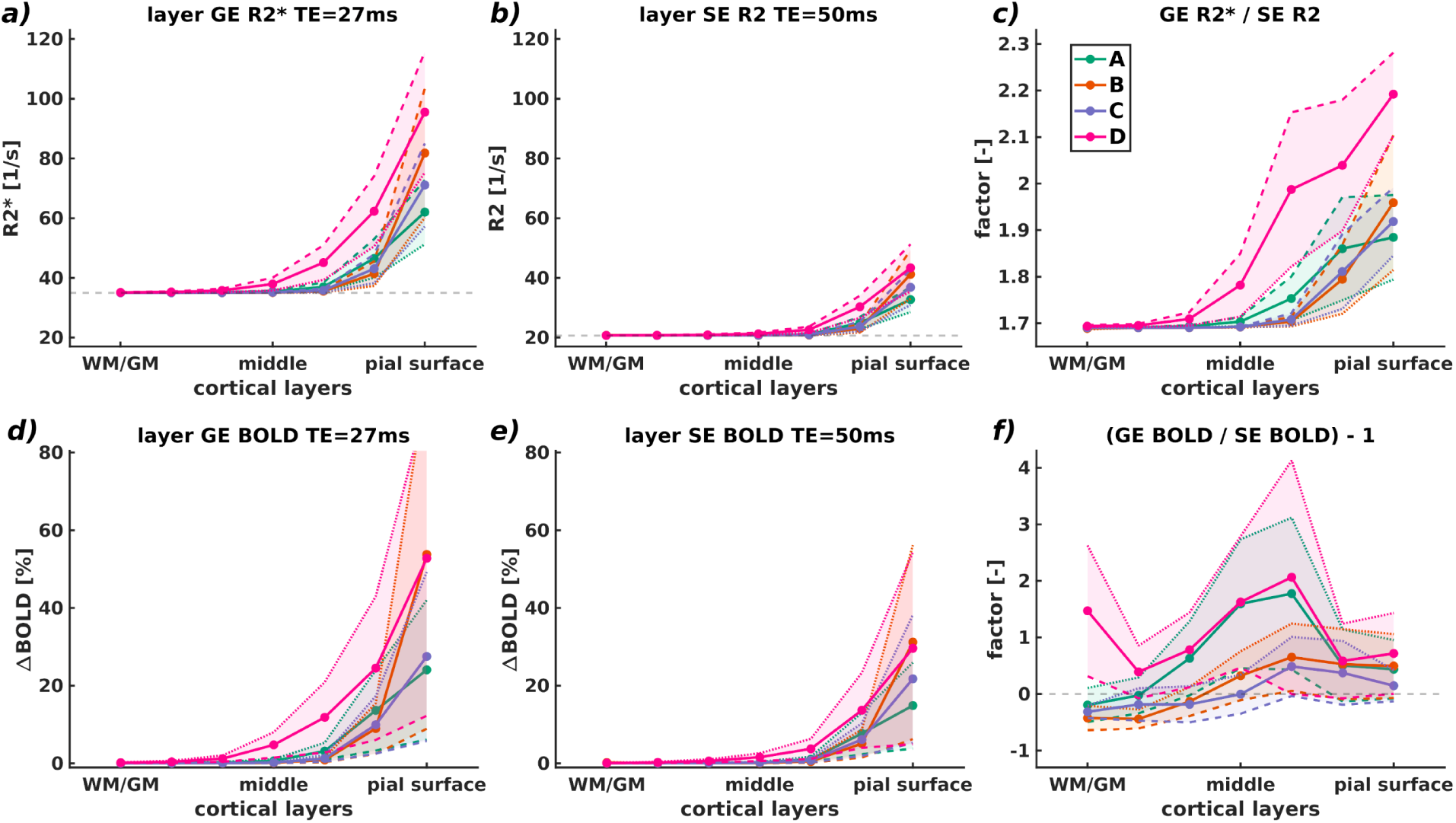
The relationship between R2* and R2 decay rates (panels **a) – c)**) and BOLD signal changes (panels **d)**-**f)**) for both GE (TE = 27 ms) and SE (TE = 50 ms) across cortical depth (seven equidistant layers) is depicted for all vascular models. Panels **c)** and **f)** show the resulting GE R2*/SE R2 ratio and the ((GE BOLD/SE BOLD) – 1) ratio, respectively. The solid lines represent mean values across all oxygen saturation levels, with shaded areas indicating 95% confidence intervals computed by Monte Carlo simulations. The dashed line indicates the boundary corresponding to the lowest oxygen saturation value, while the dotted line represents the boundary corresponding to the highest oxygen saturation value. The horizontal dashed line in panels **a)** and **b)** corresponds to the intrinsic R2_0_(*) of tissue. The bottom row panels show solid lines indicating mean BOLD signal change across all oxygen saturation levels, using 59% oxygen saturation state as reference condition (SO2vein = 59%). Shaded areas represent 95% confidence intervals. GE R2* values were larger near cortical pial surface, decreasing toward deeper layers. SE R2 exhibited similar trend to GE R2* but was reduced/damped by refocusing pulse. Notably, the refocusing pulse did not entirely eliminate the contribution of large vessels at cortical pial surface.

GE R2* decay rates are larger near the cortical pial surface and demonstrates a decreasing trend toward the deeper layers, where most of the R2* contribution comes from the tissue’s R2*_0_ (≈ 35.71 [1/s] accounting for weighted average from extravascular and intravascular CBV and small contributions from R2’). SE R2 decay rates exhibit a similar trend as GE R2* (increasing values toward the cortical pial surface) but are smaller due to the refocusing 180-degree pulse - most of the R2 contribution in deeper layers comes from the tissue’s R2_0_ (≈ 20 [1/s] accounting for weighted average from extravascular and intravascular CBV). However, the refocusing pulse do not entirely eliminate the contribution of large vessels at the pial surface. The factor between GE R2* and SE R2 in deeper layers (panel ***c)*** towards the WM/GM boundary) is approximately 1.70. Conversely, at the top layers (towards the cortical pial surface), the ratio ranges between approximately 1.90 and 2.25. While GE R2*, SE R2, and the GE R2*/SE R2 ratio are comparable across the A, B, and C models, the D vascular model exhibited different behavior, showing a pronounced difference in the GE R2*/SE R2 ratio compared to the other models.

Similarly, the BOLD signal change displayed in panels ***d)*** and ***e)*** shows higher values near the cortical pial surface (mean GE BOLD: ∼30% signal change; SE BOLD: ∼15% signal change) compared towards the deeper layers (mean GE BOLD: ∼3% signal change; SE BOLD: ∼2% signal change). The ratio between GE BOLD and SE BOLD (panel ***f)***) demonstrates a similar trend for the A and D model, and comparable ones to the B and C model - ranging between -0.5 and 2.0 depending on the cortical depth and vascular model.

At the cortical pial surface, the ((GE BOLD/SE BOLD) - 1) ratio for all vascular models (panel ***f)***) are above the reference line. At middle layers, the factor increases to approximately two-fold dependent on the vascular structure - suggesting a large contribution from microvascular and macrovascular compartments. Moreover, at deeper layers, the signal is mostly dominated by the SE readout for all vascular models except the D vascular model - suggesting contribution only from microvascular compartments.

In Figure 5, the dependence of R2^(^*^)^ decay rates across different echo times for the seven equidistant cortical layers is shown for all vascular models. Each solid line in the results represents the mean value, while the shaded area indicates the 95% confidence interval averaged across all the oxygen saturation levels. The R2^(^*^)^ decay rates of the cortical pial surface exhibit the most significant impact on the laminar MR signal compared to deeper cortical layers, for both GE and SE sequences. It is notable that SE R2 reaches a steady-state for all cortical layers at longer echo times, with its value dependent on cortical depth. Closer proximity to the pial surface corresponds to higher R2^(^*^)^ values. In contrast, GE R2* of the pial surface layer exhibits lower values for longer echo times, underscoring its dependence on the sequence parameters. Additionally, the confidence intervals for both GE R2* and SE R2 are wider for superficial layers than deeper layers. For layers deeper than the middle layer, neither GE R2* nor SE R2 demonstrate considerable changes relative to the intrinsic R2_0_^(^*^)^ of the tissue, respectively.

**Figure 5.**
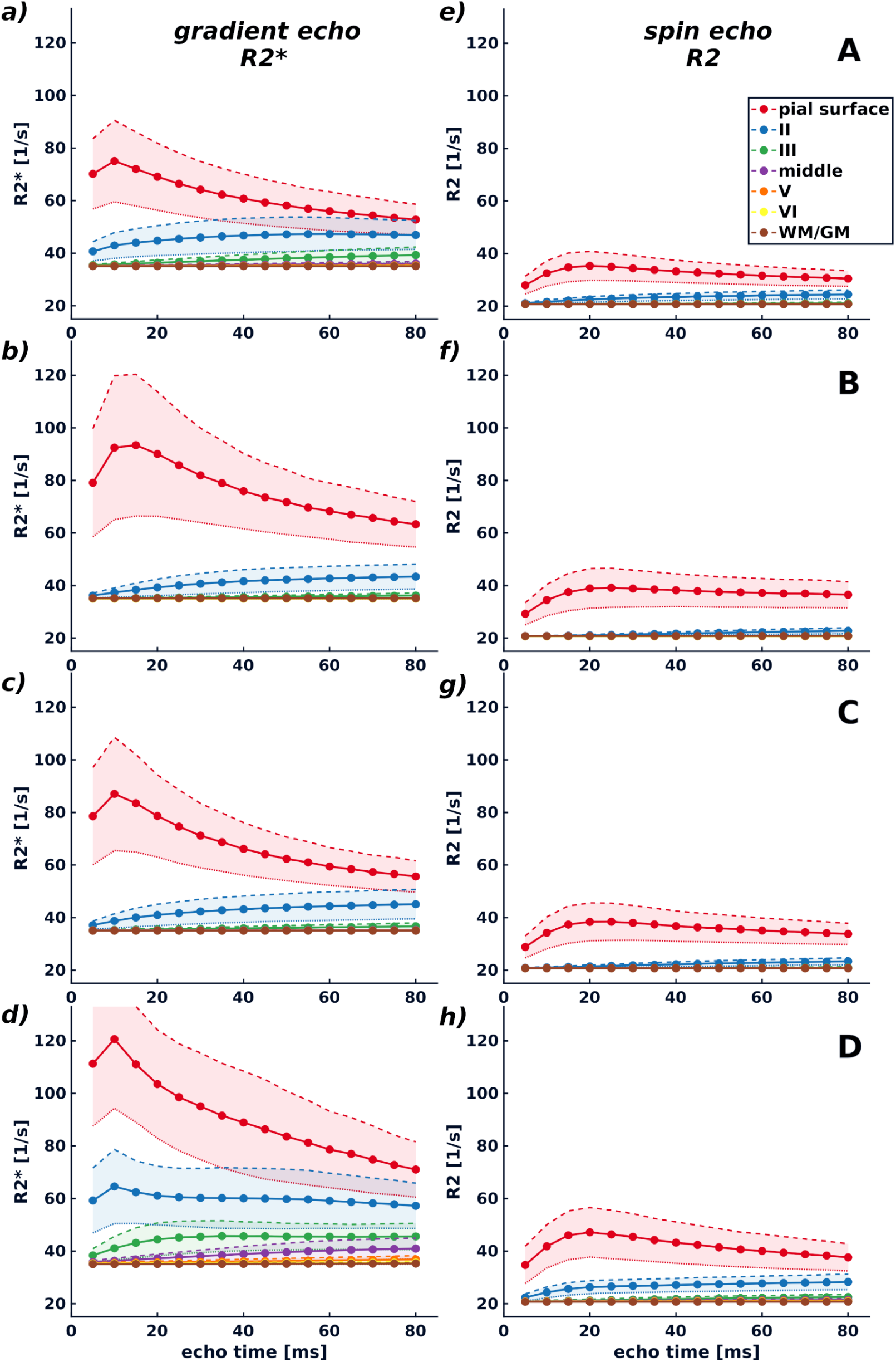
Layer GE R2* and SE R2 and its dependence on echo time. The correlation between R2^(*)^ decay rates at different echo times across seven equidistant cortical layers for all vascular models is shown. Each solid line in the graphs represents the mean value, while the shaded area indicates the 95% confidence interval using all oxygen saturation states. The decay rates of R2^(^*^)^ at the cortical pial surface have the most pronounced impact on the MR signal contribution compared to deeper cortical layers, evident in both GE and SE sequences. Consistent with previous observations, GE R2* shows larger values than SE R2. Notably, SE R2 stabilizes across all cortical layers with longer echo times, its value varying with cortical depth. Proximity to the pial surface correlates with higher GE R2* values. Conversely, GE R2* displays a slight decrease across echo times at the pial surface layer, emphasizing its reliance on the choice of echo time in sequence parameter selection. Moreover, the confidence interval for both GE R2* and SE R2 is broader for superficial layers compared to deeper layers. From the middle to deeper layers, neither GE R2* nor SE R2 exhibits substantial changes relative to the tissue’s intrinsic R2 ^(*)^.

In Figure 6, we show the BOLD signal changes for different echo times across cortical depth for all vascular models. Each solid line in the results represents the mean value, while the shaded area indicates the 95% confidence interval averaged across all the oxygen saturation levels. Notably, the cortical pial surface layer shows the most prominent BOLD signal change for both GE and SE, followed by the layer labeled with the number II and III. The rest of the deeper cortical layers (IV to WM/GM) present similar behaviors across models, except for the D model, which displays a relatively larger contribution from deeper layers. SE BOLD signal changes display comparable, to certain degree, behavior for all vascular models.

**Figure 6.**
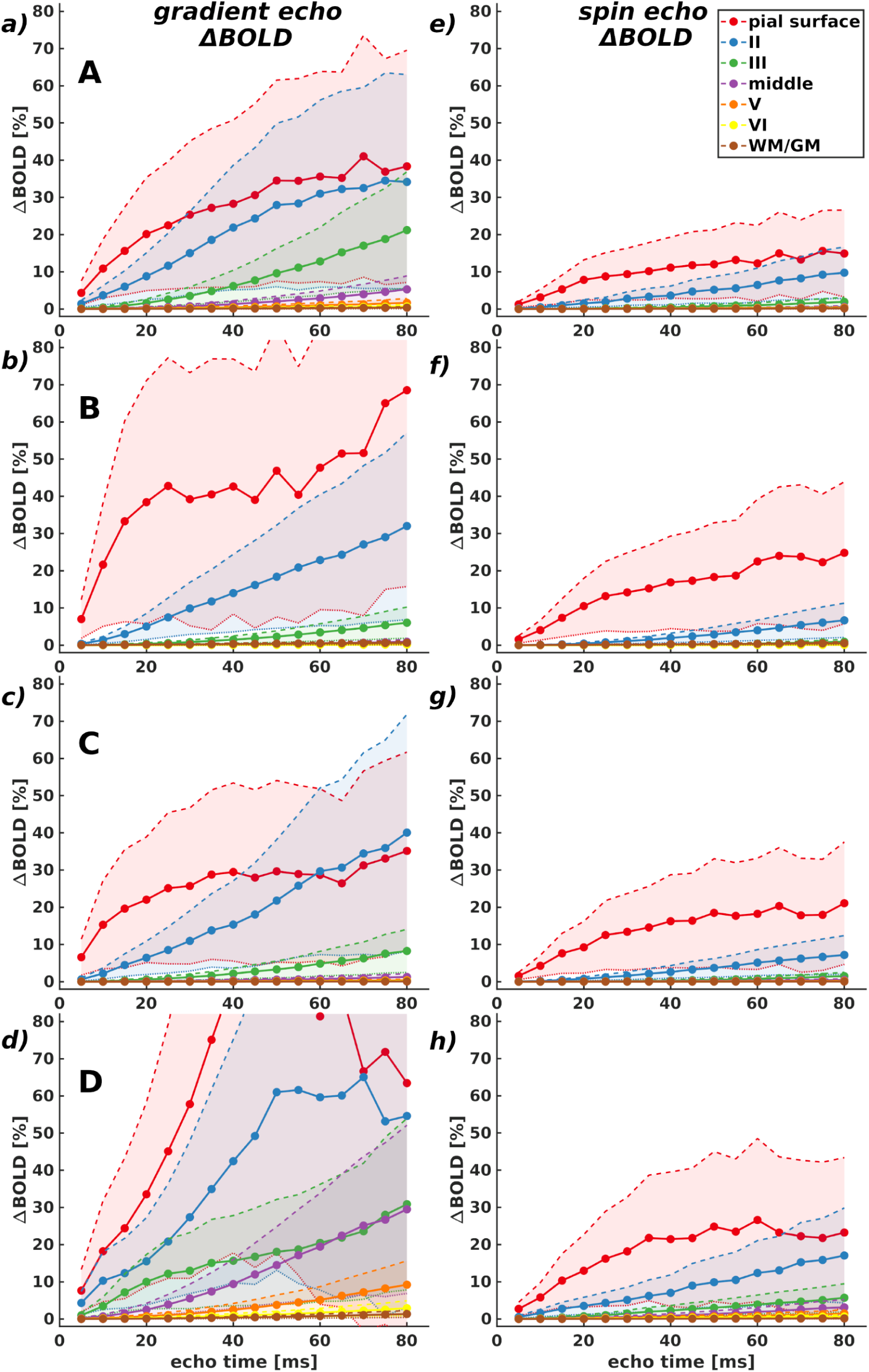
GE BOLD and SE BOLD signal changes across different echo times. The variations in BOLD signal across cortical depth and different echo times are examined for all (A - D) vascular models. Each solid line in the graphs represents the mean value, while the shaded area indicates the 95% confidence interval averaged across all the oxygen saturation levels. The cortical pial surface exhibits the most substantial BOLD signal change for both GE and SE, followed by the layer labeled with the number II and III. The remaining deeper cortical laminae (IV to WM/GM) exhibit similar behaviors across models, with the exception of the D model, which demonstrates a comparatively larger contribution from deeper layers.

## 4. DISCUSSION

### 4.1 General discussion

In this study, our aim was to characterize the contribution of diverse vascular architectures to R2^(^*^)^ and BOLD signal changes using gradient echo and spin echo readouts across cortical depths at 7T. We utilize four realistic 3D vascular models derived from two-photon imaging data and by adjusting the artery-vein ratio, we replicate the human cortical vasculature. We simulated specific oxygen saturation states and biophysical interactions and show that the specificity and signal amplitude of BOLD signals at 7T are determined by the spatial distribution of large vessels across cortical depth and on the pial surface. Our simulations indicate that spin echo BOLD imaging cannot fully mitigate the macrovascular contribution near the cortical pial surface. Additionally, we find that the intravascular contribution, stemming from both arteries and veins, is not negligible in spin echo acquisitions, particularly at high oxygen saturation levels.

Simulations using these realistic 3D vascular models reveal that R2(*) values and BOLD signal changes are influenced by extravascular and intravascular contributions across cortical depth for both GE and SE sequences. GE R2* values and BOLD signal changes exhibit an increase near the pial surface, mirrored by SE R2 decay rates and BOLD signal changes, although to a lesser extent due to the refocusing 180-degree pulse. Interestingly, the refocusing pulse did not entirely eliminate the contribution of large vessels at the pial surface, indicating that SE BOLD may not fully mitigate macrovascular effects near this cortical layer. Furthermore, our results suggest that the vascular architecture deeper in gray matter, predominantly comprising microvessels, has a relatively minor impact on the laminar signatures of both BOLD signal changes and R2(*), resulting in comparable outcomes across the four vascular models. Additionally, specific CBV values for micro- and macrovasculature vary with voxel size and position within the voxel. Hence, based on our simulation results, we conclude that the specificity and signal amplitude of BOLD fMRI signals, whether obtained using spin or gradient echo sequences, are primarily determined by the spatial distribution of large vessels across cortical depth and on the cortical pial surface

### 4.2 Diverse vascular architecture affects the BOLD fMRI signal formation

The vascular architectures used are composed of arteries, microvessels (capillaries, small arterioles/venules), and veins. The capillary bed within the four models exhibits similar vascular properties, e.g., vessel radius, relatively similar baseline CBV and vessel density across cortical depth (see Figure 1), and to a certain degree, comparable topologies. It is important to note that the vascular architectures were obtained from the same cortical region (parietal), but different mice. The microvascular similarities suggest that, despite the limited number of vascular models, the MR signal contribution from this compartment is comparable across simulations as demonstrated by the relative similar patterns in frequency shifts at the middle plane (see Figure 2). The macrovascular architecture is different in all models. This make evident that the diverse topology of the large vessels has a high impact on the MRI signal formation in a voxel, as demonstrated in the inhomogeneous frequency shifts at the top plane (see Figure 2), and thus, the contribution from large vessels might not be generalizable, particularly at high spatial resolutions.

It has been previously recognized that the initial phase of the MR signal decay diverges from a mono-exponential curve, as noted in studies such as those by Yablonskiy et al., [2010]^38^, Kiselev et al., [1999]^30^. Our simulations have validated this observation. The global R2^(^*^)^ decay values at the voxel level across the four vascular models consistently demonstrated a nonlinear response to changes in oxygen saturation for both gradient echo and spin echo readout techniques at 7T (Figure 3). However, SE R2 presents a smaller dephasing effect compared to GE R2* due to the effect of the 180-degree radiofrequency refocusing pulse. The variations in the computed R2^(^*^)^ behavior concerning oxygen saturation among the models can be attributed to differences in their vascular structure. As mentioned earlier, significant differences in the topology of the macrovasculature are observed, while the microvasculature shows almost comparable features across all vascular models which results in similar local inhomogeneous magnetic fields. Simulations using the mono-sized randomly placed and oriented cylinder models^24^ show that large vessel contribution is still present in the SE BOLD signal change but reduced as compared with the GE BOLD signal change within the plausible physiological oxygen saturation range as shown in the ***Supplementary*** Figure 2.

### 4.3 The laminar BOLD fMRI signal is dependent on pulse sequence selection and its imaging parameters

GE R2* decay rates show an increase towards the cortical pial surface. SE R2 decay rates follow a similar pattern to GE R2*, increasing towards the cortical pial surface, albeit to a lesser extent due to the refocusing 180-degree pulse. However, even with the refocusing pulse, the influence of large vessels at the cortical pial surface remains significant. In the deeper layers, most of the contributions to GE R2* and SE R2 originate from the tissue’s R2_0_^(*)^, accounting a small weighted contribution from extravascular and intravascular CBV and small contributions from R2’. Our findings suggest that the diverse vascular architecture in deeper gray matter has a diminished effect on the laminar signatures of both BOLD signal changes and R2^(^*^)^ decay rates. Specifically, the variability in the amplitude of BOLD signal changes and R2^(^*^)^ decay rates is reduced in deeper layers, enabling more reliable comparisons of laminar data across individuals or conditions. Nonetheless, minor differences persist even in deeper layers. On the other hand, superficial layers (pial surface) exhibit significant differences in the topology of the large vessels, leading to a less uniform fingerprint BOLD signal change in these layers. However, these simulation results support the necessity of addressing the bias of large vessels toward the pial surface in laminar fMRI data, for example, through filtering and/or normalization techniques^51,56^.

The choice of echo time in both pulse sequences impacts the contribution of BOLD signals across cortical depth. Overall, our results suggest that the largest BOLD signal contribution at different echo times is observed in the cortical layers near the superficial layer, i.e the pial surface. However, this contribution depends on the topology of the vascular architecture, as middle layers can also play a significant role in generating the BOLD contrast at particular echo times.

### 4.4 Methodological considerations

The BOLD signal is affected by alterations in local CBF, CBV, and CMRO_2_. In this study, we have focused solely on simulating the effects of various oxygen saturation states. However, it is important to incorporate changes in cerebral blood volume in future investigations, as variations in CBV significantly influence R2(*) decay values and, consequently, the BOLD signal change^13,38,46,47^.

Furthermore, this encourages deeper exploration of the complex relationship between neural activity, hemodynamic response, and the unique characteristics of cortical layers. While our study introduced a more realistic vascular architectural model to assess its impact on the BOLD signal across cortical depth, we noted the relative simplicity of our hemodynamic model. Acknowledging this limitation, we recognize the necessity of adopting a more sophisticated hemodynamic model for future advancements. Advancing our understanding of hemodynamic processes will refine simulations by incorporating factors like blood flow dynamics, oxygen transport, and metabolic regulation, leading to a more comprehensive representation of vascular function and its influence on imaging signals^45,55^.

The calculation of the BOLD signal change in this study focuses on the extravascular and intravascular signal contribution. The extravascular component arises from the interaction between diffusing spins and magnetic field changes due to susceptibility differences in the vascular compartments and due to differential oxygen saturation levels. The intravenous contribution to the BOLD signal might be disregarded for GE readouts due to the relatively short T2*_0_ relaxation time constant of oxygenated blood at ultra-high magnetic fields, as indicated by Uludağ et. al. [2009]^29^. While the intravascular contribution of both arterial and venous components may be negligible for GE sequences, the intravascular component of the arterial compartment is demonstrated to play an important role in SE sequences at 7T.

It has been demonstrated that the specificity of the SE acquisition scheme has a clear dependence on the echo train length used to encode the signal using EPI readouts^31,32^. In this work we presumed signal acquisition without any influence of the imaging gradients –duration of spatial position encoding is neglected.

Thus, different parameter selection in the EPI readouts may influence the behavior of the R2^(^*^)^ and the corresponding BOLD signal changes^32^.

The vascular specificity of SE R2 also depends on the assumed diffusion coefficient. The diffusion coefficient at the GM/CSF boundary (superficial pial layer) might influence the computed BOLD signal changes due to the faster diffusional motion of water molecules in the CSF compartment. Future simulations should take into account these two diffusion motion regimes^38,48^.

The fluctuation in the amplitude of the BOLD signal concerning the angular orientation of the main magnetic field is significantly influenced by the orientation of the vasculature, as supported by previous studies in human subjects^12,54^ and simulations^45,58^. This angular dependency is directly attributed to both pial vessels and penetrating/ascending vessels. Therefore, we envision future studies to demonstrate the laminar variability in the amplitude of the BOLD signal depending on various oxygen saturation levels.

A limitation of the current study is the arbitrary labeling of the large vessels. Here, we enforced an assumed human artery/vein ratio for labeling the macrovessels. While this ratio was consistent across all models, the specific selection of certain vessels might influence the behavior of the laminar profile across cortical depth.

Likewise, it’s crucial to consider the disparity in cortical thickness between mice and humans^39,45,53^, though this does not influence the simulation conclusions reported here. Nevertheless, we anticipate improving our computational model to account for this disparity. One prospective method could entail virtually generating synthetic 3D vascular networks^45^, incorporating statistical characteristics of both macrovessels and microvessels, which span approximately ∼2-3 mm in cortical depth and maintain an artery-venous ratio consistent with the cortical properties of a specific brain region^53^.

### 4.5 Conclusion

We observe that GE R2* decay rates follow an increasing trend towards the cortical pial surface, peaking near this layer, and demonstrate a decreasing trend towards the deeper layers. In the deeper layers, the majority of the R2(*) contribution originates from the tissue’s intrinsic R2(*)0. SE R2 decay rates exhibit a similar pattern to GE R2* with increasing values towards the cortical pial surface; however, they are smaller due to the presence of the refocusing 180-degree pulse. Despite the inclusion of the refocusing pulse, the contribution of large vessels at the cortical pial surface remains significant. This pattern is similar for the BOLD signal changes. Therefore, the diverse topology of large vessels influences the relative sensitivity of the MRI signal to different layers, and the reliance on specific models suggests that contributions from large vessels may not be generalizable. Furthermore, while the vascular compartments remain consistent across the four vascular models, variations in vascular topology, particularly in the macrovasculature, result in differences in computed R2(*) effects and BOLD signal changes. Thus, the use of multiple realistic 3D vascular models is necessary in studies exploring the relationship between angioarchitecture and the biophysical effects on the human fingerprint BOLD fMRI signal formation. This computational approach helps in understanding the influence of the vascular architecture on GE BOLD and SE BOLD signal formation across cortical depth, as well as the impact of pulse sequence parameter selection on BOLD signal changes at submillimeter acquisitions.

## ACKNOWLEDGEMENTS

This work was supported by the National Institute of Mental Health of the National Institutes of Health under the Award Number R01MH111417 and the Dutch Research Council under award number 18969. The content is solely the responsibility of the authors and does not necessarily represent the official views of the National Institutes of Health.

## AUTHOR CONTRIBUTION STATEMENT

MGBY, JS and NP: conceived and designed the project. NP and MvO: obtained funding to support the project. MGBY: developed the computational pipeline, run the simulations, designed the figures and wrote the original draft of the manuscript. VC: performed the labelling of the vessels and analyzed the vascular data. All authors discussed the results and reviewed and edited the manuscript.

## DISCLOSURE/CONFLICT OF INTEREST

The authors declare that they have no known competing financial interests, conflict of interest or personal relationships that could have appeared to influence the work reported in this paper.

## CODE/DATA AVAILABILITY STATEMENT

The code and data underlying the findings of this study are available from the corresponding author upon request. Access is subject to a nonexclusive, revocable, non-transferable, and limited right to use solely for research and evaluation purposes, excluding any commercial use.

## ABBREVIATIONS

3D: three-dimensional
7T: 7 tesla
BOLD: blood oxygenation level-dependent
CBF: cerebral blood flow
CBV: cerebral blood volume
CMRO2: oxygen metabolism
CSF: cerebrospinal fluid
fMRI: functional magnetic resonance imaging
GE: gradient echo
GM: grey matter
HcT: hematocrit
SE: spin echo
TE: echo time
WM: white matter

## Notes

### Competing Interest Statement

The authors have declared no competing interest.

