## Supplemental Figures for "On the influence of the vascular architecture on Gradient Echo and Spin Echo BOLD fMRI signals across cortical depth: a simulation approach based on realistic 3D vascular networks"

*Supplementary Figures*

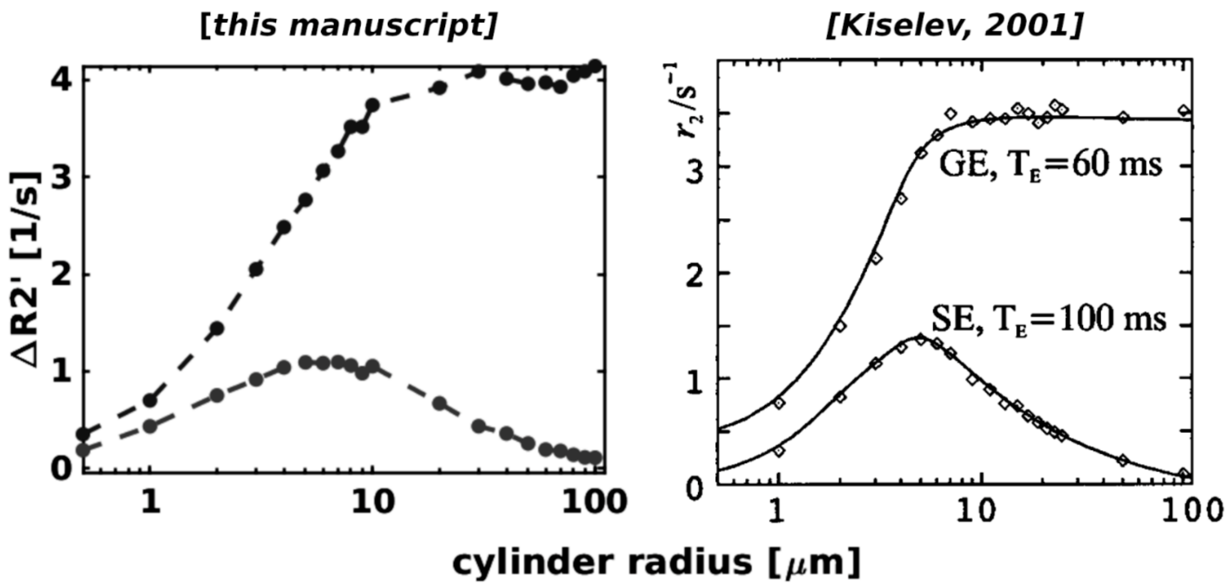


***Supplementary Figure 1****. A classic result regarding vessel type contribution was reproduced. The ΔR2' decay rate (left panel) was computed to validate our Monte Carlo simulation pipeline, as presented in [Kiselev, 2001] (right panel). This ΔR2' reproduction assumes diffusion effects and an imposed susceptibility difference (Δχ = 1E-7). This simulation was performed for both GE (TE = 60 ms) and SE (TE = 100 ms) at 1.5T.*

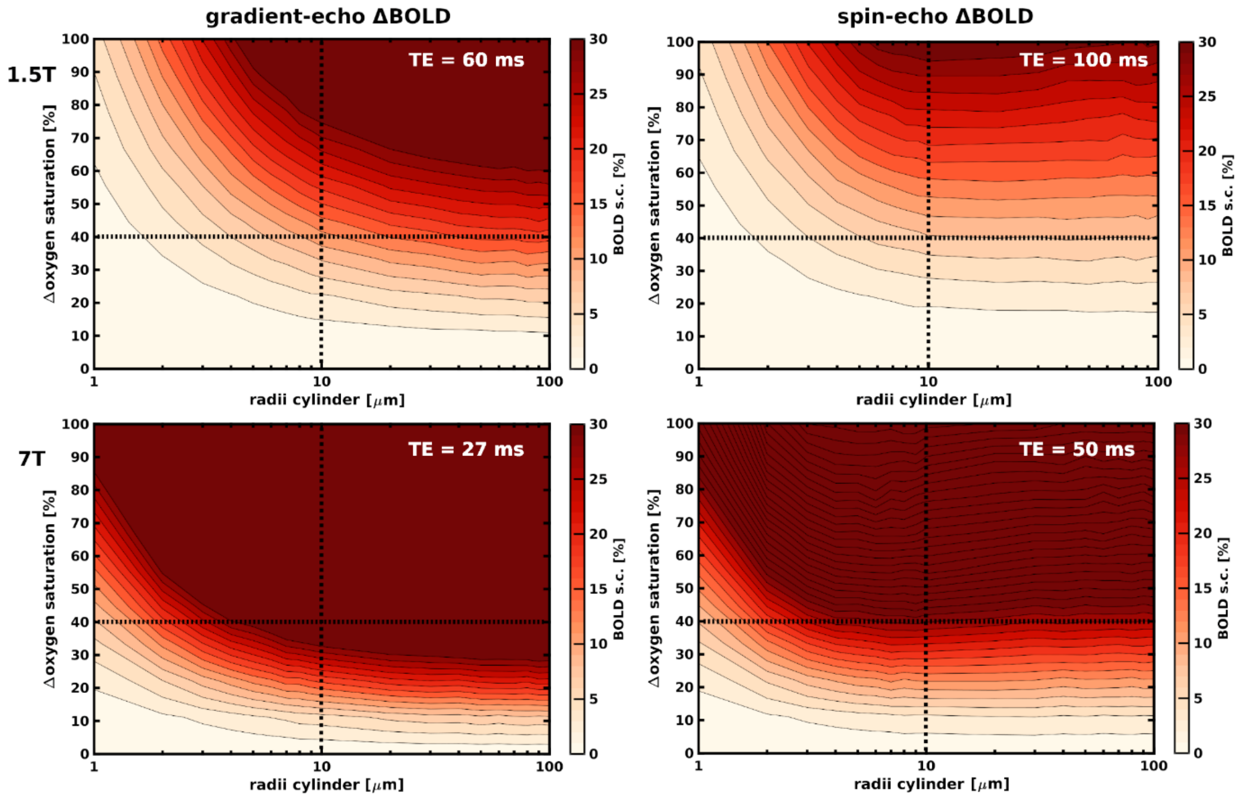


***Supplementary Figure 2**. Simulated BOLD signal changes using randomly oriented monosize cylinder voxel models for GE and SE at 1.5T (top row) and 7T (bottom row) using their respective echo times as indicated on the graphs. The vertical black dotted lines represent the assumed separation between the macrovascular (large radii values) and microvasculature (small radii values) contributions to the BOLD signal change. Furthermore, we assumed that the values below the horizontal black dotted lines represent physiologically plausible values capable of generating these BOLD signal changes. Assuming the baseline state as fully oxygen-saturated (1.0), the selected value of 40% oxygen saturation corresponds to an oxygen saturation of 0.60. The parameters of the simulation are similar as described in [Kiselev et al., 2001] –D = 1E-9 $m^2$/s, volume fraction = 3%, HcT = 40%, time-step = 50 μs.*

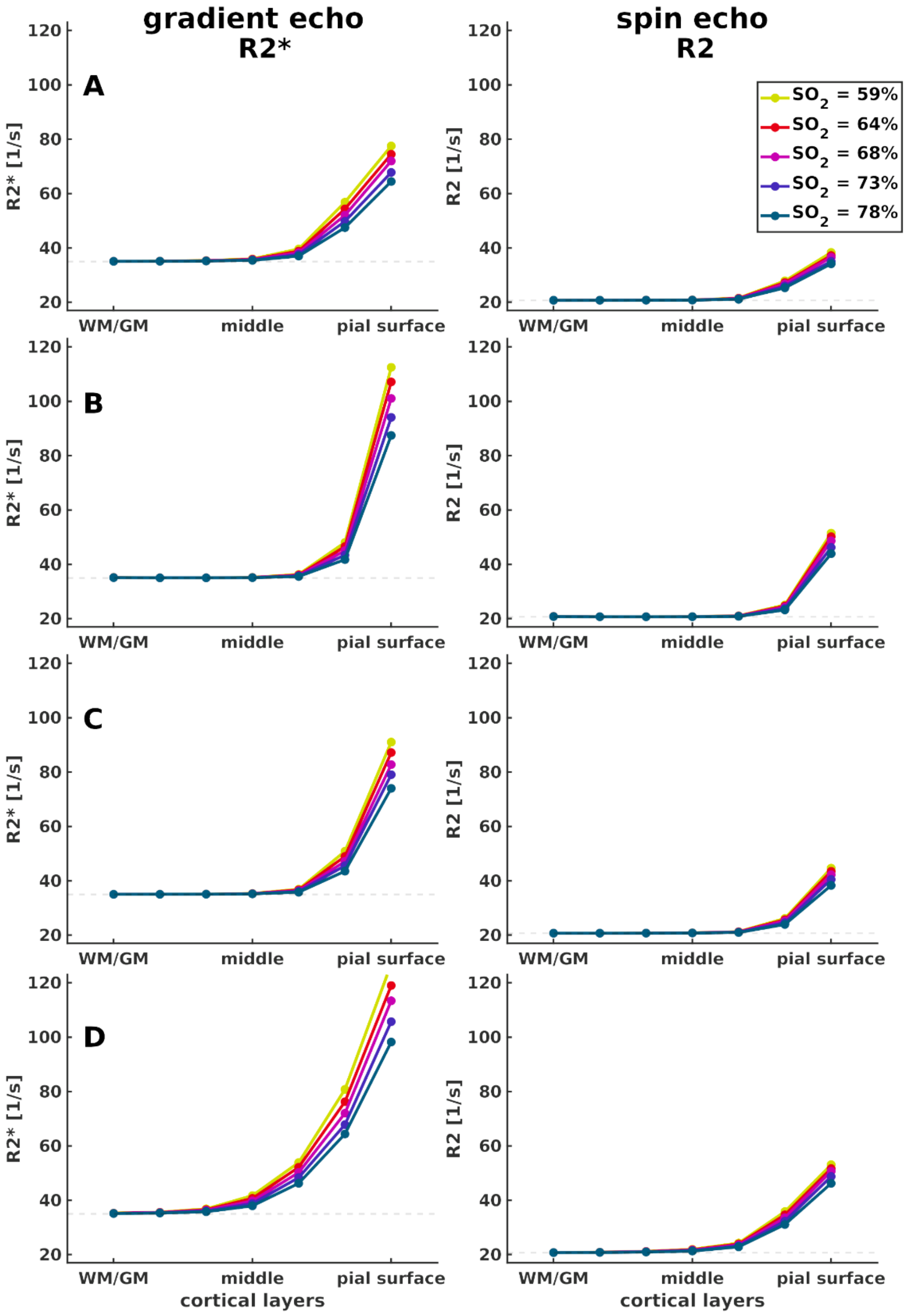


***Supplementary Figure 3a***. *The relationship between R2* and R2 decay rates for both GE (TE = 27 ms) and SE (TE = 50 ms) across cortical depth (seven equidistant layers) is depicted for all vascular models (A –D) for all physiological plausible oxygen saturation levels (59% to 78%) independently. This result is used to generate the plots in* ***Figure 3a*** *and* ***3b***.

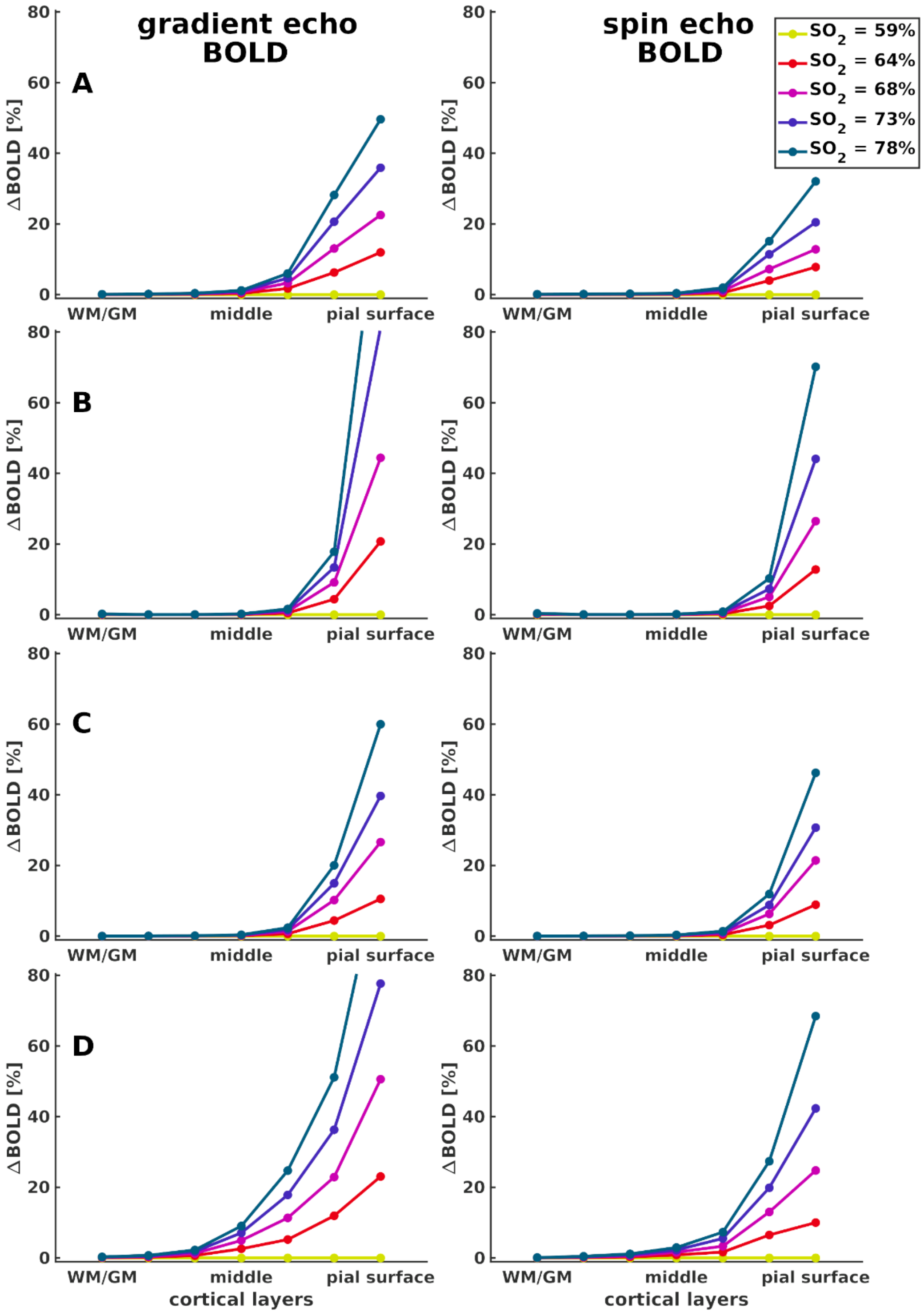


***Supplementary Figure 3b***. *The relationship between BOLD signal changes for both GE (TE = 27 ms) and SE (TE = 50 ms) across cortical depth (seven equidistant layers) is depicted for all vascular models (A -D) for all physiological plausible oxygen saturation levels (59% to 78%) independently. This result is used to generate the plots in* ***Figure 3d*** *and* ***3e***.
